# Activin receptor type IIA/B blockade increases muscle mass and strength, but compromises glycemic control in mice

**DOI:** 10.1101/2025.06.11.659035

**Authors:** Michala Carlsson, Emma Frank, Joan M. Màrmol, Mona Sadek Ali, Steffen H. Raun, Edmund Battey, Nicoline Resen Andersen, Andrea Irazoki, Camilla Lund, Carlos Henríquez-Olguin, Martina Hubec Højfeldt, Pauline Blomquist, Frederik Duch Brome, Andreas Lodberg, Christian Brix Folsted Andersen, Marco Eijken, Jonas Roland Knudsen, Erik A. Richter, Lykke Sylow

## Abstract

**Purpose:** Blocking the Activin receptor type IIA and B (ActRIIA/IIB) has clinical potential to increase muscle mass and improve glycemic control in obesity, cancer, and aging. However, the impact of blocking ActRIIA/IIB on strength, metabolic regulation and insulin action remains unclear.

**Methods:** Here, we investigated the effect of short- (10 mg/kg once, 40h) or long-term (10 mg/kg twice weekly, 21 days) antibody targeting ActRIIA/IIB (αActRIIA/IIBab) in lean and diet-induced obese mice and engineered human muscle tissue.

**Results:** Short-term *α*ActRIIA/IIB administration in lean mice increased insulin-stimulated glucose uptake in skeletal muscle by 76-105%. Despite this, *α*ActRIIA/IIB-treated mice exhibited 33% elevated fasting blood glucose and glucose intolerance. Moreover, long-term *α*ActRIIA/IIB treatment increased average muscle mass (20%) and reduced fat mass (-8%) in obese mice but did not change insulin-stimulated glucose uptake in skeletal muscle or adipose tissue, yet induced marked glucose intolerance, and increased hepatic glucose output in response to pyruvate. Concomitantly, long-term *α*ActRIIA/IIBab treatment increased strength (30%) in mouse soleus muscle and prevented activin A-induced loss of tissue strength in engineered human muscle tissue. Surprisingly, long-term *α*ActRIIA/IIBab treatment lowered volitional running (-250%).

**Conclusion:** Our findings demonstrate that, in accordance with human studies, ActRIIA/IIB blockade holds promise for increasing muscle mass, strength, and insulin sensitivity. However, contrary to the improved glycemic control in humans, ActRIIA/IIB blockade in mice causes severe glucose intolerance and lowers voluntary physical activity. Our study underscores the complex metabolic and functional consequences of ActRIIA/IIB blockade, and highlight species differences on glycemic control, which warrant further investigation.

## 1. Introduction

Maintaining muscle mass and glycemic control is vital for human health. However, muscle loss occurs with pharmacologically induced weight loss and is a hallmark of chronic conditions such as cancer, aging, obesity, and type 2 diabetes, contributing to reduced survival ^1^. Muscle accounts for up to 75% of insulin-stimulated glucose disposal in humans and is therefore crucial metabolic regulation ^2–6^. Thus, preserving muscle mass and glycemic balance is vital. Despite extensive preclinical and clinical efforts, no approved therapies address both.

Emerging targets to improve both muscle function and glucose homeostasis are the transforming growth factor-β (TGF-β) family ligands, such as activin A and myostatin ^7^. These ligands inhibit skeletal muscle growth via the activin receptor type IIA and type IIB (ActRIIA/IIB) ^8,9^. Conversely, the endogenous activin A and myostatin inhibitor, follistatin, increases muscle mass ^10–13^, and improves muscle insulin sensitivity towards glucose uptake and protein synthesis in mice^1^. Accordingly, blocking activin A and myostatin promotes hypertrophy and prevents muscle loss, as seen in mice with cancer-induced weight loss^14^ and in mitigating semaglutide-induced muscle loss in obese mice ^15^. Yet, understanding the potential ActRIIA/IIB-regulated link between muscle mass regulation and insulin-mediated glucose uptake remains incomprehensible. Moreover, exploring therapeutic interventions addressing muscle metabolism is promising for managing conditions associated with concomitant muscle wasting and insulin resistance ^16,17^.

Bimagrumab is a human monoclonal antibody that blocks the ActRIIA/IIB ^18–21^, and has been evaluated in Phase II trials^22,23^. An increased glucose infusion rate during a hyperinsulinemic-euglycemic clamp and reduced HbA1c in insulin-resistant individuals^19,23^ suggest that Bimagrumab enhances insulin sensitivity, which is corroborated by a recent study in primates showing reduced HbA1c levels upon myostatin and activin A blockade on top of GLP1-RA treatment^24^ However, in mice, emerging evidence suggests that ActRIIA/IIB blockade may disrupt glucose homeostasis, given that hepatic follistatin production caused glucose intolerance ^25^ and Bimagrumab treatment elevated blood glucose levels ^15^.

Considering the growing aging population prone to develop obesity and conditions associated with muscle wasting, such as sarcopenia, cancer, and diabetes, investigating the effect of ActRIIA/IIB blockade on muscle mass and strength, insulin-mediated glucose uptake, and glycemic control is crucial.

Here, we present data showing that while ActRIIA/IIB blockade holds promise for enhancing muscle mass, function, and insulin sensitivity, it also induces severe glucose intolerance and reduces voluntary physical activity in mice, underscoring the complex metabolic and functional consequences of ActRIIA/IIB blockade and highlighting the importance delineating species differences with these potential therapeutics.

## 2. Methods

### 2.1 Animals

All experiments were approved by the Danish Animal Experimental Inspectorate (License: 2021-15-0201-01085). Male C57BL/6JRj mice (Janvier lab, France), were maintained on a 12 h:12 h light-dark cycle and single-housed at thermoneutral temperature (29 °C), with nesting and hiding material. A 10 weeks diet intervention was initiated at 14 weeks of age, where lean mice received a standard rodent chow diet 3.1 kcal/g (Altromin no. 1324; Brogaarden, Hørsholm, Denmark) and tap water, and diet-induced obese (DIO) mice received a 45% high fat 4.75 kcal/g (Research diet no. D12451 2.5 HS; Brogaarden, Hørsholm, Denmark) and 10% sucrose water *ad libitum*.

### 2.2 Body composition analysis

Changes in lean mass and fat mass were determined by quantitative magnetic resonance imaging (MRI) using an Echo MRI scanner (EchoMRI-4in1TM, Echo Medical System LLC, Texas, USA).

### 2.3 αActRIIA/IIBab treatment

αActRIIA/IIBab (αActRIIA/IIB) was purified from conditioned media using chromatography and exchanged into phosphate – buffered saline (phosphate buffered saline, PBS). The amino acid sequence of the anti-ActRIIA/IIB antibody was identical to the commercial variant known as Bimagrumab. More details about the production and purification of αActRIIA/IIB can be found in ^26^. Mice were intraperitoneally (i.p) treated short-term (10mg/kg bw, 40h) or long-term (10mg/kg bw), two times pr week for 21 days with the αActRIIA/IIB.

### 2.4 Glucose tolerance test

Before the test, the mice were fasted for 4 hours. D-mono-glucose (2g kg^-1^ bw) was administered i.p., and blood was collected from the tail vein. Blood glucose was determined at 0, 20, 40-, 60-, 90-, and 120-minutes post glucose injection using a glucometer (Bayer Contour, Bayer, Switzerland). Plasma from 0 and 20 minutes was collected, frozen in liquid nitrogen. Insulin levels were measured in duplicates. (Mouse Ultrasensitive Insulin ELISA, #80-INSMSU-E01ALPCO Diagnostics, USA).

### 2.5 Pyruvate tolerance test

The mice were fasted for 4 hours before sodium pyruvate (1 g kg^-1^ bw) was administered i.p and blood collected from the tail vein. Blood glucose was determined at time points 0, 20, 40, 75, and 90 minutes using a glucometer (Bayer Contour, Bayer, Switzerland).

### 2.6 *Ex vivo* force assessment

Mice were euthanized by cervical dislocation and soleus muscles were rapidly isolated and non-absorbable 4–0 silk suture loops (Look SP116, Surgical Specialities Corporation) were attached at both ends. The muscles were placed in a DMT Myograph system (820MS; Danish Myo Technology, Hinnerup, Denmark) and incubated at 30°C in a Krebs-Ringer buffer and electrical stimulation protocol was performed as previously described ^27^.

### 2.7 Insulin tolerance test and *in vivo* 2-deoxy glucose measurements

To determine whole-body insulin tolerance and 2-deoxy-glucose (2DG) uptake in muscle, [3H]2DG (Perkin Elmer) was injected retro-orbitally in a bolus of saline containing 66.7 μCi mL^-1^ [3H]2DG (6 μLg^-1^ bw) in lean and DIO mice, as previously described ^28^.

### 2.8 Protein extraction and immunoblotting

All tissues were processed, and lysate protein concentration was determined using the bicinchoninic acid method, and immunoblotting was performed as previously described ^29^.

### 2.9 RNA extraction and real-time-qPCR

RNA from mouse livers was extracted using the RNeasy Mini Kit (Qiagen, #74106) following the manufacturer’s instructions as previously described^27^. Primer Sequence: *PCX*_Forward: CTGAAGTTCCAAACAGTTCGAGG; *PCX*_Reverse: CGCACGAAACACTCGGATG; *PCK1*_Forward: CTGCATAACGGTCTGGACTTC; *PCK1*_Reverse: CAGCAACTGCCCGTACTCC

### 2.10 Cross-sectional area analysis

Cross-sectional area (CSA) was assessed as described in ^30^. Cryosections were thawed, air-dried, rehydrated, and blocked with 5% goat serum before overnight incubation at 4°C with rabbit anti-Laminin (Merck, L9393; 1:150) followed by a 1-hour incubation at room temperature (RT) with Alexa Fluor 488-conjugated goat anti-rabbit (Invitrogen, A-11008; 1:800). After washing, sections were mounted with Prolong Gold Antifade and imaged using a Zeiss Axioscan.Z1. CSA was quantified by thresholding the Laminin signal in FIJI ^31^.

### 2.11 Insulin-stimulated glucose uptake human myotubes

Primary myoblasts from Cook Myosite healthy donor 30M were seeded in a 96-well plate (16,000 cells/well) in Myotonic growth media containing MS-3333 MyoTonic^TM^ Growth Supplement and 1% P/S. Following two days of growth, the media was changed to MyoTonic^TM^ differentiation media (MD-5555). On day 4 of differentiation, cells were treated with 1 µM αActRIIA/IIB (Cat #: HY-P99355, MedChem) for 96h. Insulin-stimulated glucose uptake was assessed on day 7 using the Glucose Uptake-Glo Assay (Promega, Cat. #J1343) following the manufacturer’s recommendations. In brief, on the day of the experiment, the human myotubes were starved three hours before insulin stimulation. The cells were stimulated with insulin (Novo Nordisk) for 60 min at 37°C, 5% CO_2_. Insulin solution was removed and 50 µL 2DG (10 mM) was added for 10 minutes and placed on a shaker at RT. Following 2DG transport, 25 µL stop buffer was added and placed on a shaker for 10 min at RT. Then, 25 µL neutralization buffer was added, and the plate was shaken briefly. The 2DG6P detection reagent was added to the plate and incubated for 60 min at RT. After incubation, luminescence was recorded using a ClarioStar Plus (BMG Labtech, Germany).

### 2.13 Generation of engineered muscle tissue from primary human myoblast

Engineered muscle tissues were generated as previously described^32^. In brief, primary myoblasts from Cook Myosite healthy donor 30M (300,000 cells) were resuspended in 42.8µl Hams F10 medium (Gibco), supplemented with 12µl Matrigel, 4µl 50mg/mL fibrinogen (Sigma-Aldrich), and 1.2µl 100U/µl thrombin (Sigma-Aldrich) for a total seeding volume of 60µl, 5 million cells per mL. The cell suspension was applied to a Mantarray casting well with mini-sized wells and a two-post array, and incubated at 37°C, 5% CO2 for 80 min for hydrogel formation. Next, 1 mL of F10 media was added to the wells and, after an additional 10 minutes incubation, the posts were transferred to Myotonic media (Cook Myosite) supplemented with 5g/mL aminocaproic acid (Sigma-Aldrich). 24 hours after casting the engineered muscle tissues, they were transferred to Curi Bio Primary Skeletal Muscle Differentiation media and media was refreshed ever 2-3 days. After 7 days, the engineered muscle tissues were transferred to Curi-Bio Primary Skeletal Muscle Maintenance Media. At day 7, the electrical stimulations began.

### 2.14 Force assessment in engineered muscle tissue

Force production in the engineered muscle tissues was assessed every 2-3 days followed by media change. Assessment was performed using a purpose-built plate lid supporting graphite electrodes compatible with the Mantarray hardware. Force was monitored during electrical stimulation at 1-2-3-5-1020-30 and 40Hz each at a 2sec duration with 8 seconds in between stimulations. Each stimulation consisted of biphasic 75mAmp pulses of 10ms duration^32^.

### 2.15 Statistical analysis

Results are shown as mean ± standard error of the mean (SEM) with the individual values shown for bar graphs or mean ± SE for curve graphs. Statistical testing for normally distributed data was performed using t-test, one-way or two-way (repeated measures when appropriate) ANOVA as applicable. Sidak post hoc test was performed for all ANOVAs to test the difference between control and of *α*ActRIIA/IIB treatment. Statistical analyses were performed using GraphPad Prism, version 9 (GraphPad Software, La Jolla, CA, USA, RRID: 002798). The significance level for all tests was set at α<0.05.

## 3. Results

### 3.1 Short-term ActRIIA/IIB receptor blockade improved muscle insulin sensitivity but caused whole-body glucose intolerance

The short-term effects of *α*ActRIIA/IIB on insulin sensitivity in adipose and skeletal muscle tissues are unknown. To uncover this, we treated lean mice with a single dose (10 mg/kg bw) of *α*ActRIIA/IIB, which did not influence bw (Fig. 1A), lean (Fig. 1B) or fat (Fig. 1C) mass, or food intake (Fig. 1D) 40 hours post-injection. Yet, insulin-stimulated muscle glucose uptake was nearly doubled in all muscles, except extensor digitorum longus (EDL) (Fig. 1E). We next assessed the direct effects of ActRIIA/IIB blockade on insulin-stimulated glucose uptake in primary human myotubes. ActRIIA/IIB blockade elevated glucose uptake 65% at baseline and 75% during insulin stimulation (Fig. 1F). These results show that, both in mouse skeletal muscle and human myotubes *in vitro*, short-term ActRIIA/IIB blockade enhanced insulin-stimulated glucose uptake. *α*ActRIIA/IIB’s insulin-sensitizing effect seemed selective for the muscle since adipose tissue insulin-stimulated glucose uptake was similar between ActRIIA/IIB groups (Fig. 1G). Despite the large enhancement in skeletal muscle glucose uptake, *α*ActRIIA/IIB-treated lean mice exhibited 14% elevated fed blood glucose (Fig. 1H) and impaired glucose tolerance (Fig. 1I), compared to PBS-treated control mice. The insulin tolerance test (ITT) showed a 40% upward shift in blood glucose in *α*ActRIIA/IIB-treated mice, though the incremental area under the curve (iAOC) remained unchanged due to baseline elevated glucose levels (Fig. 1J).

**Figure 1:**
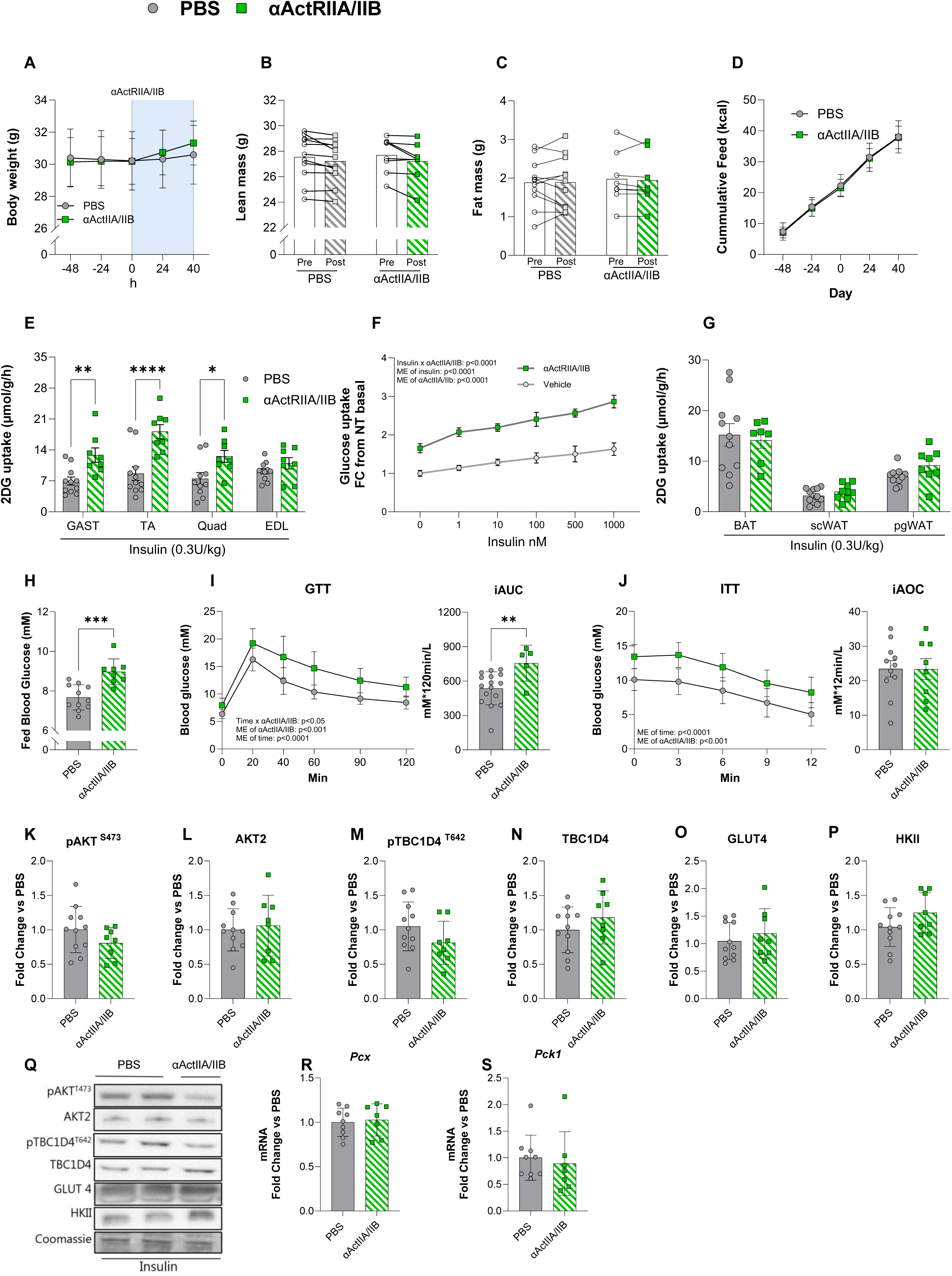
Short-term ActRIIA/IIB receptor inhibition by *α*ActRIIA/IIBab improves muscle insulin sensitivity, but causes whole-body glucose intolerance. **A)** Body weight development in control PBS treated and *α*ActRIIA/IIBab treated mice recorded 48h prior to *α*ActRIIA/IIBab treatment until 40h after treatment **B)** Magnetic Resonance Imagining-derived lean mass, and **C)** fat mass before and 40h after *α*ActRIIA/IIBab treatment. **D)** Cumulative kcal intake. **E)** 2-deoxy glucose (2DG) uptake of Gastrocnemius muscle (Gast), Tibialis Anterior (TA), Quadriceps (Quad), Extensor Digitorum Longus (EDL). **F)** Insulin-stimulated glucose uptake in primary human myotubes treated with Bimagrumab for 96h (from day 4 myotubes) to block ActRIIA/IIB signaling. Values are shown as fold change (FC) from non-treated (NT) basal. **G)** 2DG of Adipose tissue Brown adipose tissue (BAT), Subcutaneous white adipose tissue (ScWAT), and Perigonadal WAT (PgWAT). **H)** Blood glucose in the fed state 24 hours after *α*ActRIIA/IIBab treatment. **I)** Glucose tolerance test (GTT) with incremental area under the curve (iAUC) and, **J)** Insulin tolerance test (ITT) with incremental area over the curve (iAOC). Western blot-Protein expression of **K)** phosphorylated (p) pAKT^S473^cat#9271, **L)** AKTII cat#3063, **M)** pTBC1D4^T642^ cat#4288, **N)** TBC1D4 cat# ab189890, **O)** GLUT4 cat#PA1-1065, **P)** HKII cat#2867, **Q)** Representative blots from western blotting. Gene expression levels in liver tissue measured by real-time qPCR. **R)** *Pck1* liver gene expression levels S) *Pcx* liver gene expression levels. Lean PBS: n=8-16 Lean *α*ActRIIA/IIBab: n=8 (5 for GTT data). Values are shown as mean±SEM including individual values, and as mean±SD when individual values are not shown. Effect of *α*ActRIIA/IIBab or Bimagrumab: *= p<0.05, **=p<0.01, ***=p<0.001.

To elucidate the mechanisms of enhanced insulin-stimulated muscle glucose uptake, we examined key components of the insulin-signaling pathway ^33^. We observed no short-term *α*ActRIIA/IIB-induced alterations in protein content of Akt-TBC1D4 signaling (Fig. 1K, L, M, N) or glucose handling proteins GLUT4 (Fig. 1O) and hexokinase (HK) II (Fig. 1P) in gastrocnemius skeletal muscle. In addition to muscle and adipose tissue, the liver is a major regulator of glycemic control. We therefore investigated whether the elevated blood glucose in *α*ActRIIA/IIB-treated mice might be explained by increased hepatic gluconeogenesis but found no changes in *Pcx* and *Pck1* mRNA expression in liver tissue (Fig.1 R-S). Thus, short-term *α*ActRIIA/IIB treatment markedly enhanced muscle insulin-stimulated glucose uptake independent of Akt-TBC1D4 muscle insulin signaling but paradoxically elevated blood glucose and induced glucose intolerance.

### 3.2 Long-term *α*ActRIIA/IIB treatment elevated muscle mass and protected against adiposity expansion

Long-term blockade of the ActRIIA/IIB receptor has been proposed to enhance glucose homeostasis and weight loss while preventing muscle mass loss^34–36^, yet results are conflicting ^15,25^. To determine the long-term effects of *α*ActRIIA/IIB on whole-body and muscle metabolism, we treated lean and DIO mice with *α*ActRIIA/IIB for 3 weeks. Expectedly, long-term *α*ActRIIA/IIB treatment increased bw, although only in lean mice (Fig. 2A), without altered food intake (Fig. 2B). DIO PBS-treated mice displayed a 34 % increase in fat mass while *α*ActRIIA/IIB treatment blunted fat mass gain (Fig. 2C), illustrated by reduced overall fat mass and lower weights of subcutaneous white adipose tissue (-18%), perigonadal (-15%), and subscapular brown adipose tissue (-13%) compared to PBS-treated DIO mice (Fig. 2D). Lean and DIO mice treated with *α*ActRIIA/IIB displayed increased lean mass by 8% and 13%, respectively (Fig. 2E). In both lean and DIO mice, *α*ActRIIA/IIB increased muscle mass by an average 20% (Fig 2F), along with cardiac hypertrophy in DIO mice (18%). Quantification of laminin-stained gastrocnemius muscle sections revealed that *α*ActRIIA/IIB-treated lean mice displayed a 137% increase in the proportion of large fibers within the 2500–2999 µm² range compared to PBS-treated controls (Fig. 2G). Yet, this effect seemed blunted in *α*ActRIIA/IIB-treated DIO mice (Fig. 2H). Representative images are shown in Fig. 2I. These results show that long-term *α*ActRIIA/IIB treatment increased muscle mass and prevented adiposity expansion in response to DIO.

**Figure 2:**
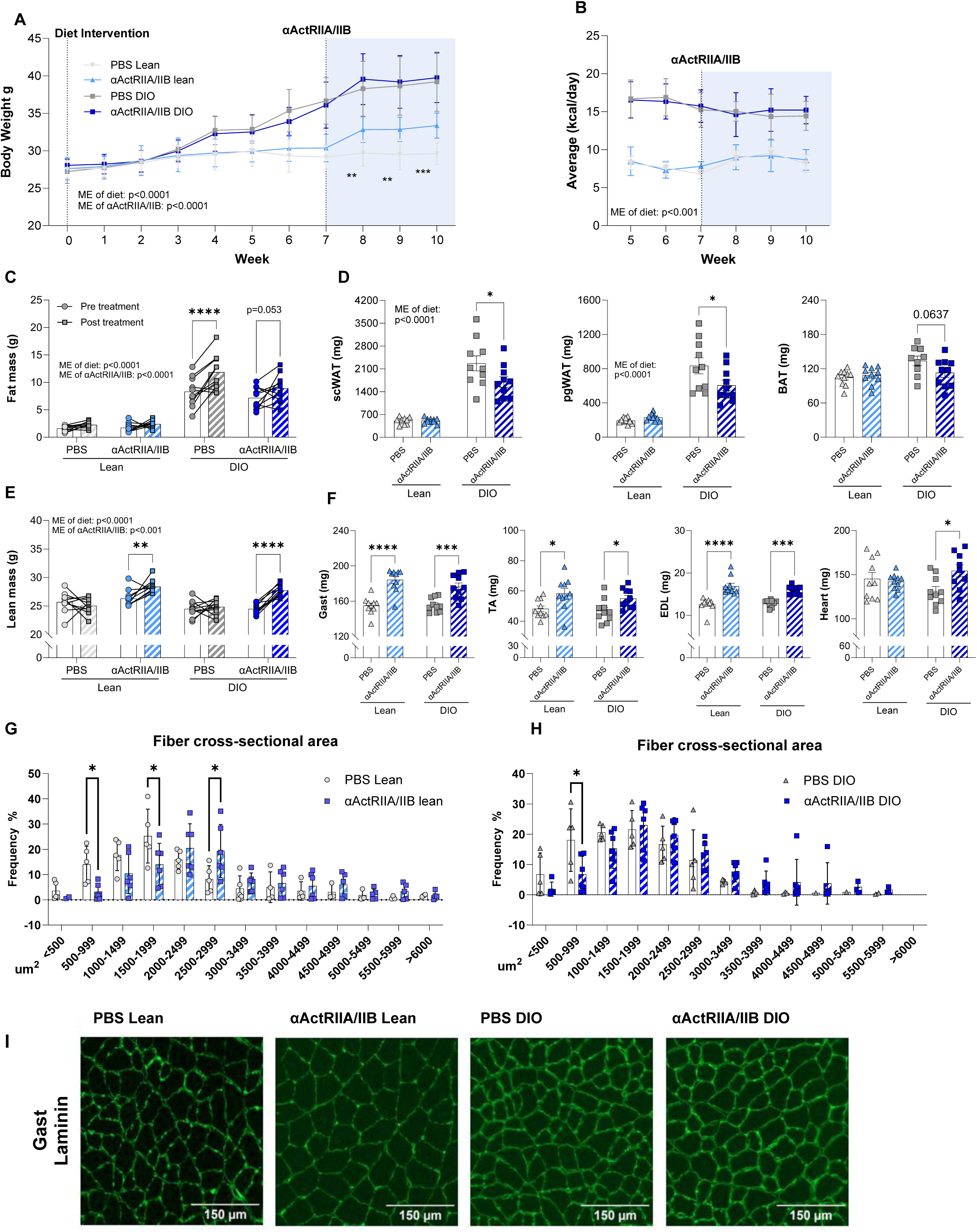
Long-term *α*ActRIIA/IIBab treatment elevated muscle mass and protected against HFHS-induced adiposity expansion. **A)** Body weight development before and after diet intervention (control chow diet (Lean) or a high fat high sucrose diet (DIO)) and *α*ActRIIA/IIBab or PBS treatment **B)** Average daily caloric intake (excluding sucrose water consumption). **C)** Magnetic Resonance Imaging-derived fat mass. **D)** Weight of subcutaneous white adipose tissue (scWAT), perigonadal WAT (pgWAT) and Brown adipose tissue (BAT). **E)** Lean mass. **F)** whole muscle weights of Gastrocnemius (Gast), Tibialis Anterior (TA), Extensor Digitorum Longus (EDL), and heart. **G)** Average fiber cross-sectional area distribution in % (single fiber segmentation average 50-300 fibers pr. picture) in Gast of lean mice and **H)** DIO mice. **I)** Representative imaging of laminin-stained Gast muscle fiber slides. PBS Lean: n=10 *α*ActRIIA/IIBab Lean: n=10 DIO PBS: n=10, DIO *α*ActRIIA/IIBab: n=10. Values are shown as mean +-SEM including individual values, and as mean+-SD when individual values are not shown. Effect of *α*ActRIIA/IIBab: *= p<0.05, **=p<0.01, ***=p<0.001. Effect of *α*ActRIIA/IIBab in DIO mice: #=p<0.05, ##=p<0.01 ###=p<0.001

### 3.3 ActRIIA/IIB blockade improves tissue strength in mouse soleus and engineered muscle tissue, but lowers running activity

Having established the beneficial effects of *α*ActRIIA/IIB on muscle mass, we next investigated the effect of *α*ActRIIA/IIB on the functional properties of the muscle, which is a poorly understood but an important clinical outcome. Following long-term *α*ActRIIA/IIB treatment in lean mice, electrically induced force production was increased in the soleus muscles (Fig. 3A), for which absolute (Fig. 3B) and specific (Fig. 3C) force production were both increased 30% (specific force, p=0.08) by *α*ActRIIA/IIB. To explore the translational clinical relevance of ActRII/IIB blockade on muscle function, we examined its effects on force production in engineered muscle tissues. We treated the engineered muscle tissues with activin A (1 nM), αActRIIA/IIB (1 µM), or in conjunction for 6 days (Fig. 3D, left panel). Activin A is an endogenous ligand for ActRIIA/IIB, and since αActRIIA/II blocks the activin A binding epitope on ActRIIA/IIB, this allowed us to evaluate αActRIIA/IIB’s potential to counteract activin A-induced suppression of force production. The presence of activin A lowered force development by 11% in engineered muscle tissues, which was blocked by αActRIIA/IIB (Fig. 3D, right panel), indicating that αActRIIA/IIB improves muscle strength in human muscle by inhibiting activin A binding to ActRIIA/IIB. Thus, in both mouse skeletal muscle and engineered muscle tissues, ActRIIA/IIB blockade elicited positive effects on muscle force production, illustrating the potential for mitigating muscle mass and functional loss.

**Figure 3:**
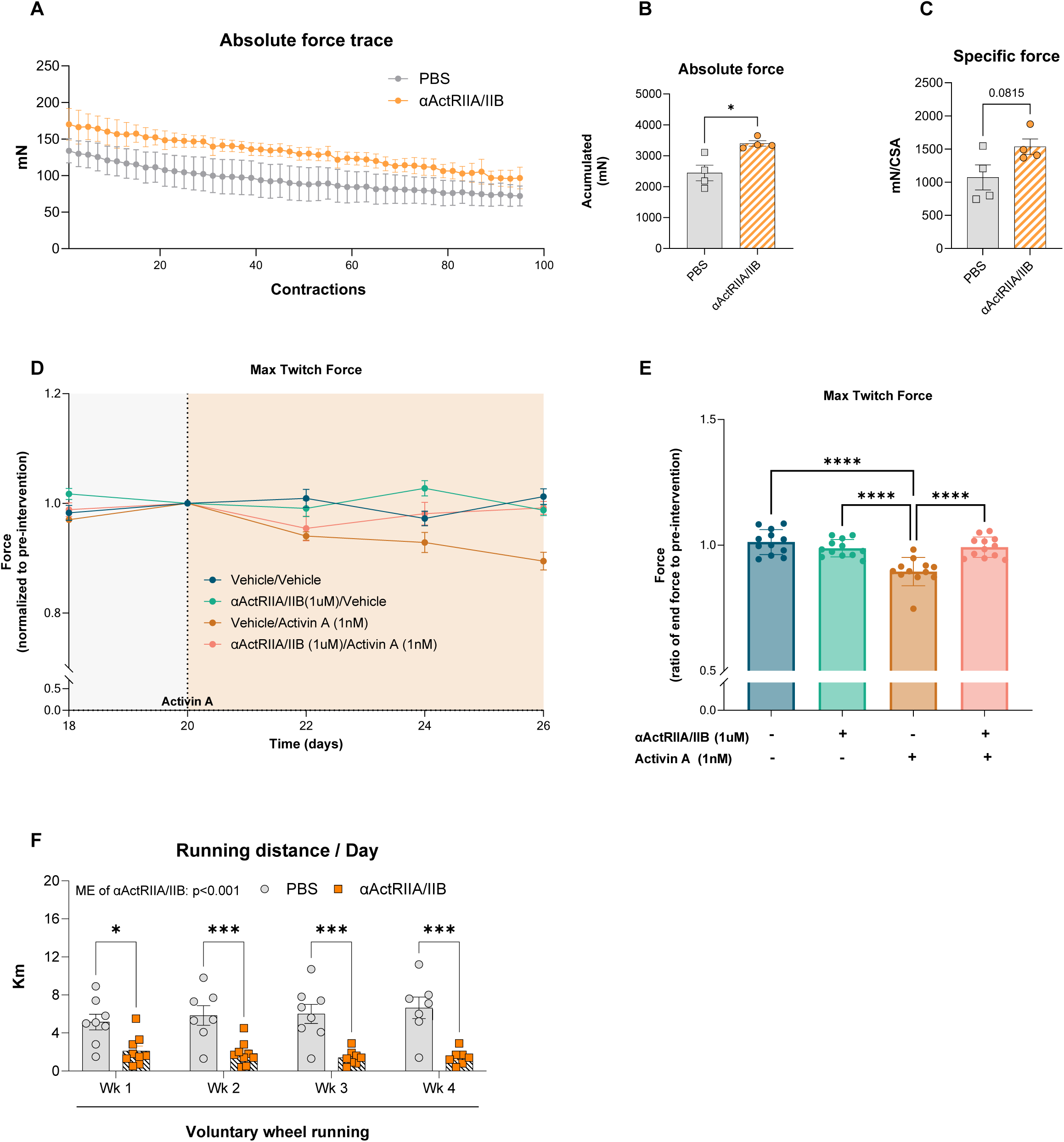
Mice treated with *α*ActRIIA/IIBab display increased muscle force ex vivo, but have markedly reduced activity in voluntary wheel running. **A)** Absolute force trace of isolated soleus muscles in *α*ActRIIA/IIBab or PBS treated mice. Soleus muscles were placed under resting tension (∼5mN) followed by electrical stimulation at 14V and 149.2 Hz with a pulse width and interval of 0.2 ms and 6.5 ms, respectively. Trains of pulses were delivered 75 times with a pause of 5000 ms between pulses, which was repeated 110 times with 2000 ms of pause between pulse trains, allowing for maximal force development. **B)** Accumulated force from absolute force trace **C)** The muscles were measured in length and weight following the electrical stimulation protocol to calculate specific force (wt(mg)/[(length(mm)*Lf*1,06]). 14-week-old male mice received two *α*ActRIIA/IIBab injections (days 0 and 14), and soleus muscle force was assessed on day 21. **D)** Force assessment in engineered human muscle tissue. Cells were treated with either vehicle or 1 µM Bimagrumab to block ActRIIA/IIB, Cat #: HY-P99355, MedChem). On day 20, activin A (1 nM) was added the respective groups. Force assessment was performed every 2 days, and force output was normalized to day 20, when activin A treatment began (left panel). Force output normalized to day 20 at the end of the intervention is shown in the right panel. Force was monitored during electrical stimulation at 1-2-3-5-10-20-30 and 40Hz each at a 2sec duration with 8 seconds in between stimulations. Each stimulation consisted of biphasic 75mAmp pulses of 10ms duration Media, including treatment groups, was changed every second day. **E)** Voluntary wheel-running intervention. 14-week-old *α*ActRIIA/IIBab treated and PBS treated mice were given free access to running wheels for four weeks. Values are shown as +-SEM including individual values, and as mean+-SD when individual values are not shown. Effect *α*ActRIIA/IIBab or Bimagrumab: *= p<0.05, ***=p<0.001, ****=p<0.0001

However, when assessing voluntary physical activity, *α*ActRIIA/IIB-treated lean mice exhibited a marked 250% reduction in voluntary wheel running (Fig. 3E), which may indicate neuromuscular changes or systemic fatigue despite increased muscle mass^37^. Yet, while the mechanisms underlying *α*ActRIIA/IIB’s activity-lowering effects remain unclear, its enhancement of muscle force production in both mouse skeletal muscle and engineered muscle tissues underscores its potential to counteract muscle mass and functional decline.

### 3.4 Long-term *α*ActRIIA/IIB treatment caused hyperglycemia and glucose intolerance without affecting insulin-stimulated glucose uptake or signaling

After establishing the beneficial effects of long-term *α*ActRIIA/IIB treatment on body composition, muscle mass, and function, we next examined its impact on glucose homeostasis in lean and DIO mice. We hypothesized that the longer-term beneficial adaptations in body composition would counter the adverse effects of *α*ActRIIA/IIB on glucose tolerance we observed in response to the short-term treatment. Yet, despite increased muscle mass and lower fat mass, long-term *α*ActRIIA/IIB treatment increased fed blood glucose levels (Fig. 4A) and induced marked glucose intolerance in lean mice and exacerbated glucose intolerance in DIO mice (Fig. 4B). This impairment was not likely attributable to defective insulin secretion, as plasma insulin levels (Fig. 4C) and islet insulin content and glucose-stimulated insulin release (not shown) were comparable between all groups.

**Figure 4:**
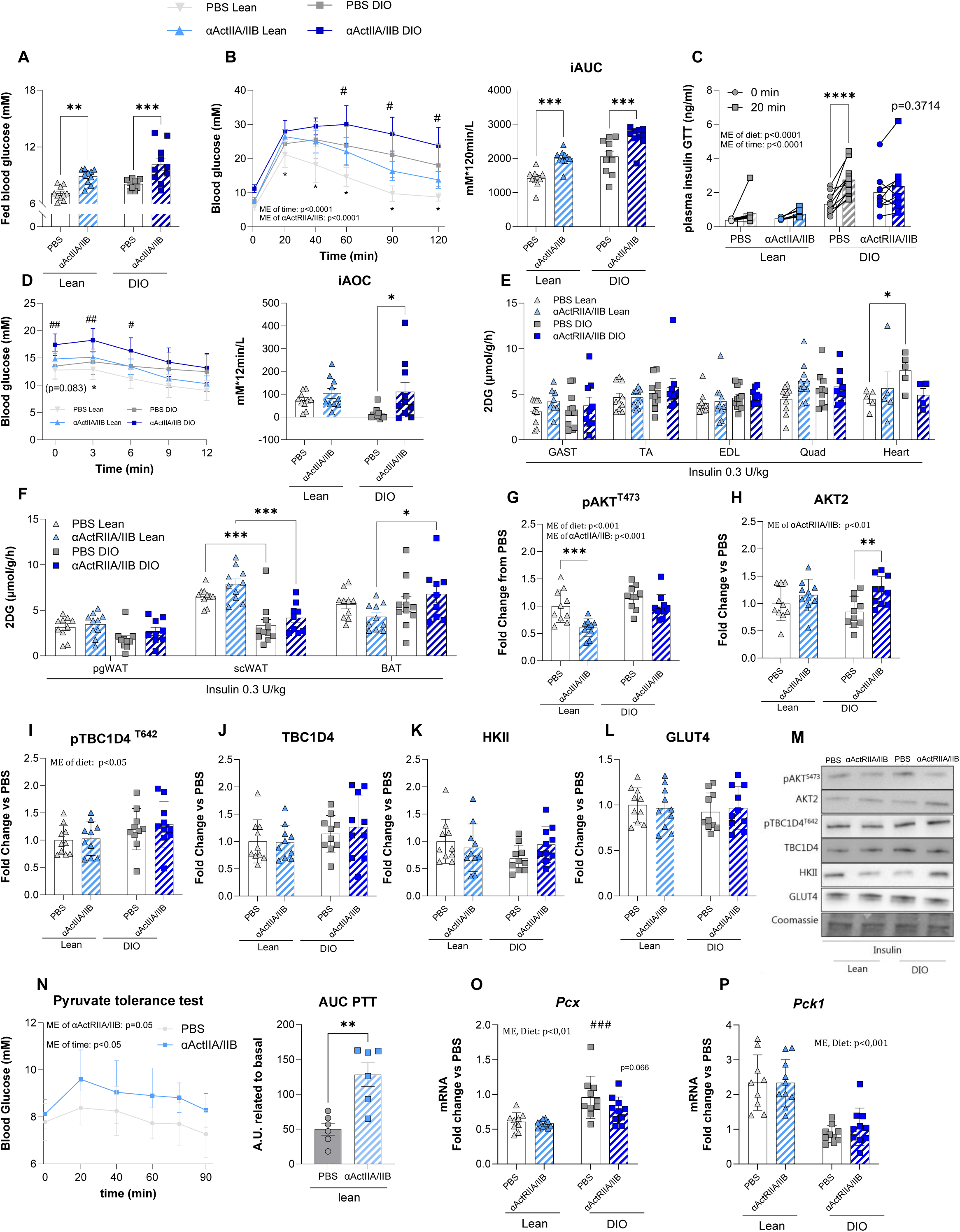
Long-term *α*ActRIIA/IIBab treatment causes whole body glucose intolerance without effect on glucose uptake and insulin signaling downstream Akt. **A)** Fed blood glucose measured the last week of long-term *α*ActRIIA/IIBab treatment. **B)** Blood glucose levels before (0min), 20min, 40 min, 60 min and 90min following an intraperitoneal glucose tolerance test (2 g/kg body weight), with incremental area under the curve (iAUC) **C)** Plasma insulin from blood drawn at timepoint 0 and after 20 min of the glucose tolerance test. **D)** Blood glucose levels measured at timepoint 0 min, 3 min, 6 min, 9 min, and 12 min following retro-orbital insulin injection (0.3 U/kg body weight) and incremental area over the curve (iAOC) during the 12 min insulin stimulation. iAUC and iAOC was calculated using the trapezoid rule. **E)** 2-deoxy glucose (2DG) uptake of Gastrocnemius muscle (Gast), Tibialis Anterior (TA), Extensor Digitorum Longus (EDL), Quadriceps (Quad), and heart. **F)** 2DG of Perigonadal white adipose tissue (PgWAT), Subcutan WAT (ScWAT), and Brown adipose tissue (BAT). Western blot-Protein expression of **G)** phosphorylated (p) pAKT^S473^cat#9271, **H)** AKTII cat#3063, **I)** HKII cat#2867, **J)** GLUT4 cat#PA1-1065, **K)** pTBC1D4^T642^ cat#4288, **L)** TBC1D4 cat# ab189890. **M)** Representative blots. **N)** Pyruvate tolerance test performed 7 days after the latest *α*ActRIIA/IIBab administration). iAUC was calculated from the basal blood glucose concentration was determined using the trapezoid rule for the 90 minutes test. Gene expression levels in liver tissue measured by real-time qPCR. O) *Pcx*, liver gene expression levels. P) *Pck1*, liver gene expression levels. Values are shown as mean +-SEM including individual values, and as mean+-SD when individual values are not shown. Effect of *α*ActRIIA/IIBab: *= p<0.05, **=p<0.01, ***=p<0.001. Effect of *α*ActRIIA/IIBab in diet induced obese mice: #=p<0.05, ##=p<0.01 ###=p<0.001

As expected, the DIO control mice exhibited insulin resistance, illustrated by a low iAOC for blood glucose in response to insulin (Fig. 4D). Four-hour fasted blood glucose was elevated in *α*ActRIIA/IIB-treated mice (Fig. 4D). Yet, the blood glucose-lowering effect of insulin relative to baseline was increased by *α*ActRIIA/IIB treatment, as indicated by iAOC being increased on average by 8-fold in DIO mice treated with *α*ActRIIA/IIB compared to DIO mice treated with PBS. As expected, insulin-stimulated glucose uptake in adipose tissue was lower in DIO mice compared to lean controls; however, skeletal muscle insulin sensitivity remained unaffected by the DIO diet. In contrast to the short-term *α*ActRIIA/IIB treatment, the long-term *α*ActRIIA/IIB treatment did not increase insulin-stimulated glucose uptake in skeletal muscle (Fig. 4E). Yet, similar to the short-term *α*ActRIIA/IIB treatment, we observed no effect of *α*ActRIIA/IIB on adipose tissue insulin-stimulated glucose uptake (Fig. 4F)

Further investigation into molecular insulin signaling revealed that *α*ActRIIA/IIB modestly reduced insulin-stimulated Akt phosphorylation on the serine site 473 (pAkt^ser473^) in lean mice (Fig. 4G), while Akt2 protein content was increased (Fig. 4H). Despite these alterations in Akt phosphorylation and total Akt2 protein content, the insulin-stimulated phosphorylation of Akt’s downstream target, TBC1D4, was unaffected by *α*ActRIIA/IIB treatment (Fig. 3I, J). Moreover, we did not observe any difference in HKII (Fig. 4K) or GLUT4 (Fig. 4L) protein content. Representative blots are shown in Fig. 4M. These data show that long-term *α*ActRIIA/IIB treatment did not affect intramyocellular TBC1D4 signaling or glucose-handling protein content, consistent with unchanged insulin-stimulated glucose uptake. Consequently, altered muscle or adipose insulin sensitivity is unlikely to underlie *α*ActRIIA/IIB-induced hyperglycemia.

Considering the liver’s key role in glycemic control, we next assessed the impact of *α*ActRIIA/IIB on hepatic glucose production using a pyruvate tolerance test. Elevated blood glucose levels in lean longterm *α*ActRIIA/IIB-treated mice throughout the test (Fig. 4N) suggest enhanced lactate-to-glucose conversion. These results could indicate an increased hepatic gluconeogenesis and glucose output, which could explain the elevated blood glucose and glucose intolerance observed in *α*ActRIIA/IIB-treated mice. However, we did not observe alterations in gluconeogenesis genes *Pcx* (Fig. 4O) and *Pck1* (Fig. 4P) that could explain this regulation. Similarly, blocking ActRII/B in streptozotocin treated diabetic mice resulted in elevated blood glucose levels compared to control mice when challenged with pyruvate.^38^

Together, these results indicate that while *α*ActRIIA/IIB enhances muscle mass and function, *α*ActRIIA/IIB also induces severe glycemic disruptions in mice, likely due to increased hepatic glucose output rather than muscle or adipose insulin resistance.

## 4. Discussion

Here, we investigated the short- and long-term effects of ActRIIA/IIB blockade on insulin sensitivity, skeletal muscle mass and function, and glucose metabolism. Our findings highlight three key outcomes. First, short-term administration of *α*ActRIIA/IIB enhanced insulin-stimulated glucose uptake in mouse and primary human muscle cells, however, this effect was not sustained with long-term *α*ActRIIA/IIB treatment. Second, ActRIIA/IIB treatment increased muscle mass and improved absolute force production in isolated murine soleus muscle and engineered muscle tissues. Third, despite these benefits, *α*ActRIIA/IIB-treated mice exhibited markedly elevated fasting blood glucose levels and glucose intolerance likely due to increased hepatic glucose output.

Our first key finding was that short-term *α*ActRIIA/IIB administration enhances insulin-stimulated glucose uptake in mouse skeletal muscle and primary human muscle cells. Our data aligns with evidence that show inhibition of ActRIIA/IIB responsive ligands such as myostatin or activin A and GDF11 *via* muscle follistatin overexpression improves insulin sensitivity^10,39^. In contrast to muscle, *α*ActRIIA/IIB did not alter adipose tissue insulin-stimulated glucose uptake in neither lean nor obese mice, suggesting that the insulin-sensitizing effect of short-term *α*ActRIIA/IIB is muscle-selective. However, the insulin-sensitizing effect of *α*ActRIIA/IIB was transient and not sustained with long-term treatment in lean or obese mice, which warrants further investigation. The molecular mechanisms by which short-term *α*ActRIIA/IIB treatment enhanced insulin sensitivity was not identified in the present study, as we found no detectable absolute changes in the Akt-TBC1D4-GLUT4 signaling pathway.

Our second key finding documents that long-term *α*ActRIIA/IIB treatment can improve muscle force production. This was evidenced in both isolated mouse skeletal muscle following *in vivo α*ActRIIA/IIB treatment, as well as engineered muscle tissues. These results indicate that blocking the ActRIIA/IIB pathway, well-documented to increase muscle mass ^15,34,40^, also has beneficial effects on muscle functional capacity, supporting previous findings of improved exercise capacity and 6-min walking test in Bimagrumab-treated mice and humans, respectively ^15,41^. Surprisingly, despite these improvements in muscle force generation, we found that *α*ActRIIA/IIB-treated mice demonstrated markedly reduced voluntary wheel running activity. Rapid hypertrophy may induce muscle stiffness or discomfort, limiting voluntary movement while still enhancing force output under forced conditions, likely due to increased muscle mass. In line with our observations, participants from clinical trials reported muscle myalgia when taking Bimagrumab ^42^.

Lastly, we were surprised by our third major finding, that despite improvements on body composition, muscle mass and function, *α*ActRIIA/IIB-treated mice exhibited markedly elevated fed- and fasting blood glucose levels, and glucose intolerance. These findings align with emerging evidence from mice studies suggesting that ActRIIA/IIB blockade may disrupt glucose homeostasis, given that hepatic follistatin production caused glucose intolerance ^25^ and *α*ActRIIA/IIB-treatment caused elevated blood glucose levels in fed mice ^15,38^, similar to our findings. However, these results are in contrast to clinical data showing reduced HbA1c in Phase II clinical trials with a single dose of Bimagrumab^19,23^, or reduced HbA1c levels upon long term myostatin and activin A inhibition in primates treated with GLP-1RA^24^. These discrepancies remain to be resolved but could be due to species-specific effects and/or differences in dosing. Given the extensive use of mice in research, determining whether divergent results arise from species differences is essential for selecting appropriate preclinical models.

In our study, the detrimental effects of *α*ActRIIA/IIB on glucose handling were not due to insulin resistance of the skeletal muscle or adipose tissue. Instead, our findings highlight a potential increase in hepatic gluconeogenesis, as indicated by enhanced lactate-to-glucose conversion in pyruvate tolerance tests, which could be a mouse-specific outcome of *α*ActRIIA/IIB treatment.

## Conclusion

Together, these results indicate that, while *α*ActRIIA/IIB enhances muscle mass and function, *α*ActRIIA/IIB also induces severe glycemic disruptions in mice, likely due to increased hepatic glucose output rather than muscle or adipose insulin resistance.

These findings underscore the need to consider the metabolic consequences of ActRIIA/IIB blockade when developing therapies for conditions such as sarcopenic obesity, cancer, and diabetes. Our results also indicate species-specific differences in response to αActRIIA/IIB, underscoring the importance of this consideration in future research.

## Acknowledgments

We thank our colleagues at the Molecular Metabolism in Cancer and Aging Group, Faculty of Health and Medical Sciences, University of Copenhagen, for fruitful discussions on this topic. We acknowledge the skilled technical assistance of Betina Bolmgren and Irene B. Nielsen (NEXS, Faculty of Science, University of Copenhagen, Denmark). We acknowledge the team at Skeletal Muscle Biology, Novo Nordisk, Måløv, especially Pia Jensen, for their assistance and expertise with cell culture experiments. We acknowledge the team of Niels Billestrup, department of Biomedical Sciences, University of Copehagen for valuable assistance with beta cell experiments. We acknowledge the Core Facility for Integrated Microscopy, Faculty of Health and Medical Sciences, University of Copenhagen. Illustrations were generated using Biorender.com (RRID:SCR_018361). Graphs were generated with GraphPad PRISM (RRID:SCR_002798).

## CRediT

Conceptualization: MC, EF, LS

Data curation: MC, EF, JMM, MSA, SHR, EB, NRA, CL, CHO, LS

Formal analysis: MC, EF

Funding acquisition: LS

Investigation: MC, EF, JMM, MSA, SHR, EB, NRA, CL, CHO, MHH, PB, FDB, AL, AB, AI, LS

Methodology: JRK, LS

Project administration: MC, JMM, EF, MSA

Resources: JRK, EAR, LS

Software: EB Supervision: LS, EAR

Validation: MC, EF

Visualization: MC, EF, LS

Writing – original draft: MC, EF, LS

Writing – review and editing: all authors

## Declaration of generative AI and AI-assisted technologies in the writing process

Statement: During the preparation of this work, the authors used ChatGPT in order to do minor corrections and shortening of sentences. After using this tool/service, the authors reviewed and edited the content as needed and take full responsibility for the content of the published article.

## References

1. Egerman, M. A. & Glass, D. J. Signaling pathways controlling skeletal muscle mass. Crit. Rev. Biochem. Mol. Biol. 49, 59–68 (2014).

2. Valenzuela, P. L., Maffiuletti, N. A., Tringali, G., De Col, A. & Sartorio, A. Obesity-associated poor muscle quality: Prevalence and association with age, sex, and body mass index. BMC Musculoskelet. Disord. 21, 1–8 (2020).

3. Tisdale, M. J. Mechanisms of cancer cachexia. Physiol. Rev. 89, 381–410 (2009).

4. Dhanapal, R., Saraswathi, T. R. & Govind Rajkumar, N. Cancer cachexia. J. Oral Maxillofac. Pathol. 15, 257 (2011).

5. Haines, M. S. et al. Association between muscle mass and insulin sensitivity independent of detrimental adipose depots in young adults with overweight/obesity. Int. J. Obes. (Lond). 44, 1851 (2020).

6. DeFronzo, R. A. & Tripathy, D. Skeletal Muscle Insulin Resistance Is the Primary Defect in Type 2 Diabetes. Diabetes Care 32, S157 (2009).

7. Zimmers, T. A. et al. Induction of cachexia in mice by systemically administered myostatin. Science 296, 1486–1488 (2002).

8. McPherron, A. C., Lawler, A. M. & Lee, S. J. Regulation of skeletal muscle mass in mice by a new TGF-β superfamily member. Nature 387, 83–90 (1997).

9. Lawlor, M. W. et al. Inhibition of Activin Receptor Type IIB Increases Strength and Lifespan in Myotubularin-Deficient Mice. Am. J. Pathol. 178, 784 (2011).

10. Han, X. et al. Mechanisms involved in follistatin-induced hypertrophy and increased insulin action in skeletal muscle. J. Cachexia. Sarcopenia Muscle 10, 1241–1257 (2019).

11. Nakatani, M. et al. Transgenic expression of a myostatin inhibitor derived from follistatin increases skeletal muscle mass and ameliorates dystrophic pathology in mdx mice. FASEB J. 22, 477–487 (2008).

12. Winbanks, C. E. et al. Follistatin-mediated skeletal muscle hypertrophy is regulated by Smad3 and mTOR independently of myostatin. J. Cell Biol. 197, 997–1008 (2012).

13. Davey, J. R., et al. Integrated expression analysis of muscle hypertrophy identifies Asb2 as a negative regulator of muscle mass. JCI insight 1, (2016).

14. Liu, C. M. et al. Myostatin antisense RNA-mediated muscle growth in normal and cancer cachexia mice. Gene Ther. 15, 155–160 (2008).

15. Nunn, E. et al. Antibody blockade of activin type II receptors preserves skeletal muscle mass and enhances fat loss during GLP-1 receptor agonism. Mol. Metab. 80, 101880 (2024).

16. Lodberg, A. Principles of the activin receptor signaling pathway and its inhibition. Cytokine Growth Factor Rev. 60, 1–17 (2021).

17. Meier, D. et al. Inhibition of the activin receptor signaling pathway: A novel intervention against osteosarcoma. Cancer Med. 10, 286 (2021).

18. Rooks, D. S. et al. Effect of bimagrumab on thigh muscle volume and composition in men with casting-induced atrophy. J. Cachexia. Sarcopenia Muscle 8, 727 (2017).

19. Garito, T. et al. Bimagrumab improves body composition and insulin sensitivity in insulin-resistant individuals. Diabetes, Obes. Metab. 20, 94–102 (2018).

20. Petricoul, O. et al. Pharmacokinetics and Pharmacodynamics of Bimagrumab (BYM338). Clin. Pharmacokinet. 62, 141–155 (2022).

21. Morvan, F. et al. Blockade of activin type II receptors with a dual anti-ActRIIA/IIB antibody is critical to promote maximal skeletal muscle hypertrophy. Proc. Natl. Acad. Sci. U. S. A. 114, 12448–12453 (2017).

22. Rooks, D. et al. Bimagrumab vs Optimized Standard of Care for Treatment of Sarcopenia in Community-Dwelling Older Adults: A Randomized Clinical Trial. JAMA Netw. Open 3, e2020836–e2020836 (2020).

23. Heymsfield, S. B. et al. Effect of Bimagrumab vs Placebo on Body Fat Mass Among Adults With Type 2 Diabetes and Obesity: A Phase 2 Randomized Clinical Trial. JAMA Netw. Open 4, e2033457–e2033457 (2021).

24. Mastaitis, J. W. et al. GDF8 and activin A blockade protects against GLP-1–induced muscle loss while enhancing fat loss in obese male mice and non-human primates. Nat. Commun. *2025 161* 16, 1–8 (2025).

25. Tao, R. et al. Inactivating hepatic follistatin alleviates hyperglycemia. Nat. Med. 24, 1058 (2018).

26. Bromer, F. D. et al. The Effect of Anti-Activin Receptor Type IIA and Type IIB Antibody on Muscle, Bone and Blood in Healthy and Osteosarcopenic Mice. J. Cachexia. Sarcopenia Muscle 16, e13718 (2025).

27. Irazoki, A. et al. Housing temperature dictates the systemic and tissue-specific molecular responses to cancer in mice. doi:10.1101/2024.05.29.596034.

28. Møller, L. L. V. et al. The Rho guanine dissociation inhibitor α inhibits skeletal muscle Rac1 activity and insulin action. Proc. Natl. Acad. Sci. U. S. A. 120, e2211041120 (2023).

29. Pham, T. C. P. et al. TNIK is a conserved regulator of glucose and lipid metabolism in obesity. Sci. Adv. 9, (2023).

30. Battey, E., Dos Santos, M., Biswas, D., Maire, P. & Sakamoto, K. Protocol for muscle fiber type and cross-sectional area analysis in cryosections of whole lower mouse hindlimbs. STAR Protoc. 5, 103424 (2024).

31. Schindelin, J. et al. Fiji: an open-source platform for biological-image analysis. Nat. Methods *2012 97* 9, 676–682 (2012).

32. Smith, A. S. T. et al. High-throughput, real-time monitoring of engineered skeletal muscle function using magnetic sensing. J. Tissue Eng. 13, (2022).

33. Sylow, L., Tokarz, V. L., Richter, E. A. & Klip, A. The many actions of insulin in skeletal muscle, the paramount tissue determining glycemia. Cell Metab. 33, 758–780 (2021).

34. Hatakeyama, S. et al. ActRII blockade protects mice from cancer cachexia and prolongs survival in the presence of anti-cancer treatments. Skelet. Muscle 6, 26 (2016).

35. Rooks, D. et al. Treatment of Sarcopenia with Bimagrumab: Results from a Phase II, Randomized, Controlled, Proof-of-Concept Study. J. Am. Geriatr. Soc. 65, 1988–1995 (2017).

36. Sidis, Y. et al. Biological activity of follistatin isoforms and follistatin-like-3 is dependent on differential cell surface binding and specificity for activin, myostatin, and bone morphogenetic proteins. Endocrinology 147, 3586–3597 (2006).

37. Hanna, M. G. et al. Safety and efficacy of intravenous bimagrumab in inclusion body myositis (RESILIENT): a randomised, double-blind, placebo-controlled phase 2b trial. Lancet Neurol. 18, 834–844 (2019).

38. Wang, Q., Guo, T., Portas, J. & McPherron, A. C. A Soluble Activin Receptor Type IIB Does Not Improve Blood Glucose in Streptozotocin-Treated Mice. Int. J. Biol. Sci. 11, 199 (2015).

39. Camporez, J. P. G. et al. Anti-myostatin antibody increases muscle mass and strength and improves insulin sensitivity in old mice. Proc. Natl. Acad. Sci. U. S. A. 113, 2212–2217 (2016).

40. Rooks, D. et al. Treatment of Sarcopenia with Bimagrumab: Results from a Phase II, Randomized, Controlled, Proof-of-Concept Study. J. Am. Geriatr. Soc. (2017) doi:10.1111/jgs.14927.

41. Spitz, R. W. et al. Blocking the activin IIB receptor with bimagrumab (BYM338) increases walking performance: A meta-analysis. Geriatr. Gerontol. Int. 21, 939–943 (2021).

42. Rooks, D. et al. Safety and pharmacokinetics of bimagrumab in healthy older and obese adults with body composition changes in the older cohort. J. Cachexia. Sarcopenia Muscle 11, 1525 (2020).

